# The TIM22 complex regulates mitochondrial one-carbon metabolism by mediating the import of Sideroflexins

**DOI:** 10.1101/2020.02.06.937920

**Authors:** Thomas D. Jackson, Daniella Hock, Catherine S. Palmer, Yilin Kang, Kenji M. Fujihara, Nicholas J. Clemons, David R. Thorburn, David A. Stroud, Diana Stojanovski

**Affiliations:** Department of Biochemistry and Molecular Biology and The Bio21 Molecular Science and Biotechnology Institute, The University of Melbourne, Parkville, Victoria, 3010, Australia; Division of Cancer Research, Peter MacCallum Cancer Centre, Melbourne, Victoria 3000, Australia; Sir Peter MacCallum Department of Oncology, The University of Melbourne, Parkville, Victoria, 3010, Australia; Murdoch Children’s Research Institute, Royal Children’s Hospital, Melbourne, Victoria, 3052, Australia; Department of Paediatrics, University of Melbourne, Melbourne 3052, Australia; Victorian Clinical Genetic Services, Royal Children’s Hospital, Melbourne, Victoria, 3052, Australia

## Abstract

The Acylglycerol Kinase (AGK) is a mitochondrial lipid kinase that contributes to protein biogenesis as a subunit of the TIM22 complex at the inner mitochondrial membrane. Mutations in *AGK* cause Sengers syndrome, an autosomal recessive condition characterized by congenital cataracts, hypertrophic cardiomyopathy, skeletal myopathy and lactic acidosis. We undertook proteomic profiling of Sengers patient fibroblasts and an AGK_KO_ cell line to map the proteomic changes that ensue upon AGK dysfunction. This uncovered extensive remodelling of mitochondrial one-carbon metabolism enzymes and showed that inner membrane serine transporters, Sideroflexins (SFXNs), are novel substrates of the TIM22 complex. Deletion of *SFXN1* recapitulates the remodelling of one-carbon metabolism observed in Sengers patient cells. Proliferation of cells lacking AGK is perturbed in the absence of exogenous serine and rescuable through addition of formate, highlighting the dysregulation of one carbon metabolism as a key molecular feature in the biology of Sengers syndrome.

## Introduction

Mitochondria perform a diverse array of functions in mammalian cells, including production of ATP, induction of apoptosis, and calcium buffering (Anderson et al., 2019). Dysfunction of mitochondria is associated with many pathologies, including cancer, diabetes, and neurodegenerative disease. Mitochondrial diseases are genetic disorders that arise due to defective ATP production within the mitochondrion (Frazier et al., 2019; Jackson et al., 2018). Mitochondria require >1500 nuclear-encoded proteins to function; these proteins are delivered to specific mitochondrial sub-compartments (outer membrane; intermembrane space, inner membrane and matrix) by translocation and sorting machineries (Wiedemann and Pfanner, 2017). Mutations in genes encoding various subunits of mitochondrial translocation machineries have been linked to a number of distinct mitochondrial diseases (Jackson et al., 2018).

The TIM22 complex is an inner membrane translocase that mediates the insertion of multipass transmembrane proteins into the mitochondrial inner membrane (Rehling et al., 2003). Its main substrates are members of the SLC25A family of metabolite carrier proteins, which possess 6 transmembrane domains (Palmieri, 2013). The TIM22 complex also mediates membrane insertion of “TIM substrates”, Tim17, Tim23 and Tim22, which possess 4 transmembrane domains (Káldi et al., 1998; Kurz et al., 1999). The TIM22 complex has been extensively studied in yeast, however recent analyses in human cells has revealed substantial divergence of the complex in higher eukaryotes. The human TIM22 complex consists of: (i) Tim22, the core pore-forming subunit; (ii) the intermembrane space chaperones Tim9, Tim10, and Tim10b; and (iii) Tim29 (Callegari et al., 2016; Kang et al., 2016) and AGK (Kang et al., 2017; Vukotic et al., 2017), which are metazoan-specific subunits of the complex.

Mutations in *AGK* cause Sengers syndrome, a severe mitochondrial disease characterised by congenital cataracts, hypertrophic cardiomyopathy, exercise intolerance and lactic acidosis (Calvo et al., 2012; Mayr et al., 2012). As well contributing to protein import at the TIM22 complex, AGK also functions as a lipid kinase, able to phosphorylate monoacylglycerol and diacylglycerol to produce phosphatidic acid and lysophosphatidic acid, respectively (Bektas et al., 2005; Waggoner et al., 2004). The lipid kinase activity of AGK is dispensable for its function at the TIM22 complex (Kang et al., 2017; Vukotic et al., 2017). Despite advances in the understanding of AGK function, how the protein’s dysfunction contributes to the molecular pathogenesis underlying Sengers syndrome is unclear.

Using proteomic profiling we set out to identify which cellular pathways are dysregulated in Sengers syndrome, with a view to defining the basis of the mitochondrial dysfunction in this disease. We mapped the mitochondrial proteomes of fibroblasts from two Sengers syndrome patients (Calvo et al., 2012; Kang et al., 2017) and identified downregulation of key enzymes involved in mitochondrial one-carbon (1C) metabolism, which generates one-carbon units for use in synthesis of metabolites, including nucleotides, amino acids and lipids (Ducker and Rabinowitz, 2017). Central to this pathway is the entry of serine into mitochondria where it is converted to glycine and formate (Ducker and Rabinowitz, 2017). By analysing the proteomic data from patient cells alongside the proteomic footprint from an AGK_KO_ HEK293 cell line, we identified sideroflexins (SFXNs), including the serine transporter SFXN1 (Kory et al., 2018), as novel substrates of the TIM22 complex. *In vitro* import analyses of SFXN proteins confirmed the requirement of the TIM22 complex for import, and we determined that loss of AGK in HEK293 cells led to dependency on exogenous serine for normal proliferation. These data suggest that modulating 1C metabolism pathways/intermediates represent a viable treatment approach for Sengers syndrome and other mitochondrial diseases where the pathway is altered.

## Results

### Mitochondrial 1C metabolism is remodelled in Sengers syndrome

Calvo et al., (2012) described two unrelated patients with mutations in *AGK*. Patient 41 (referred to as Patient 1 in this study) possessed a compound heterozygous nonsense variant (p.Y390X) and splice variant that caused a shortened transcript with a premature stop codon (c.297+2T>C, pK75QfsX12). Patient 42 (referred to as Patient 2 in this study), possessed a homozygous splice variant that caused a shortened-transcript with a premature stop codon (c.1131+1G>T, p.S350EfsX19). In pursuit of a global understanding of the mitochondrial defects occurring in Sengers syndrome, we performed label-free quantitative mass spectrometry on mitochondria isolated from: (i) AGK_KO_ HEK293 cell line (Kang et al., 2017); and (ii) patient 1 and 2 fibroblasts and 3 control fibroblasts (Calvo et al., 2012; Kang et al., 2017) (**Table 1**).

AGK_KO_ HEK293 mitochondria had substantially reduced levels of the SLC25A family (10 members beyond the 1.5x down-regulation cut off) (**Figure 1A, right panel; Figure 1D; Table 1**), confirming the central role that AGK plays in the biogenesis of mitochondrial carrier proteins. Mitochondria from patient 1 and 2 fibroblasts showed a general reduction in the levels of SLC25A proteins (**Figure 1B-C, right panels; Figure 1E; Table 1**), albeit not as significantly as in the AGK_KO_ HEK293 cell line. Interestingly, key proteins in the mitochondrial arm of one-carbon metabolism (SFXN1, SHMT2, MTHFD2, MTHFD1L) (Ducker and Rabinowitz, 2017) were downregulated in both patient fibroblast cell lines (**Figure 1B-C, right panels; Figure 1G**), suggesting that dysregulation of this pathway may occur in severe forms of Sengers syndrome. 1C metabolism generates one-carbon units required to synthesise many critical metabolites, including nucleotides, amino acids, and lipids (Ducker and Rabinowitz, 2017). The pathway has cytosolic and mitochondrial branches, and a key step is the is the entry of serine into mitochondria. Transport of serine across the inner mitochondrial membrane was recently shown to be mediated by the SFXN protein family, in particular SFXN1 (Kory et al., 2018). In line with this, SFXN protein levels were reduced in both patient 1 and patient 2 fibroblasts and AGK_KO_ HEK293 cells (**Figure 1-C, right panels; Figure 1F-G**). The SFXN family consists of five members (SFXN1-5), which are each predicted to contain five transmembrane domains embedded within the inner mitochondrial membrane (Kory et al., 2018). Based on this localisation and topology,we hypothesised that SFXNs represent a novel class of TIM22 complex substrates, and that their perturbed import in Sengers syndrome leads to dysregulation of 1C metabolism.

**Figure 1.**
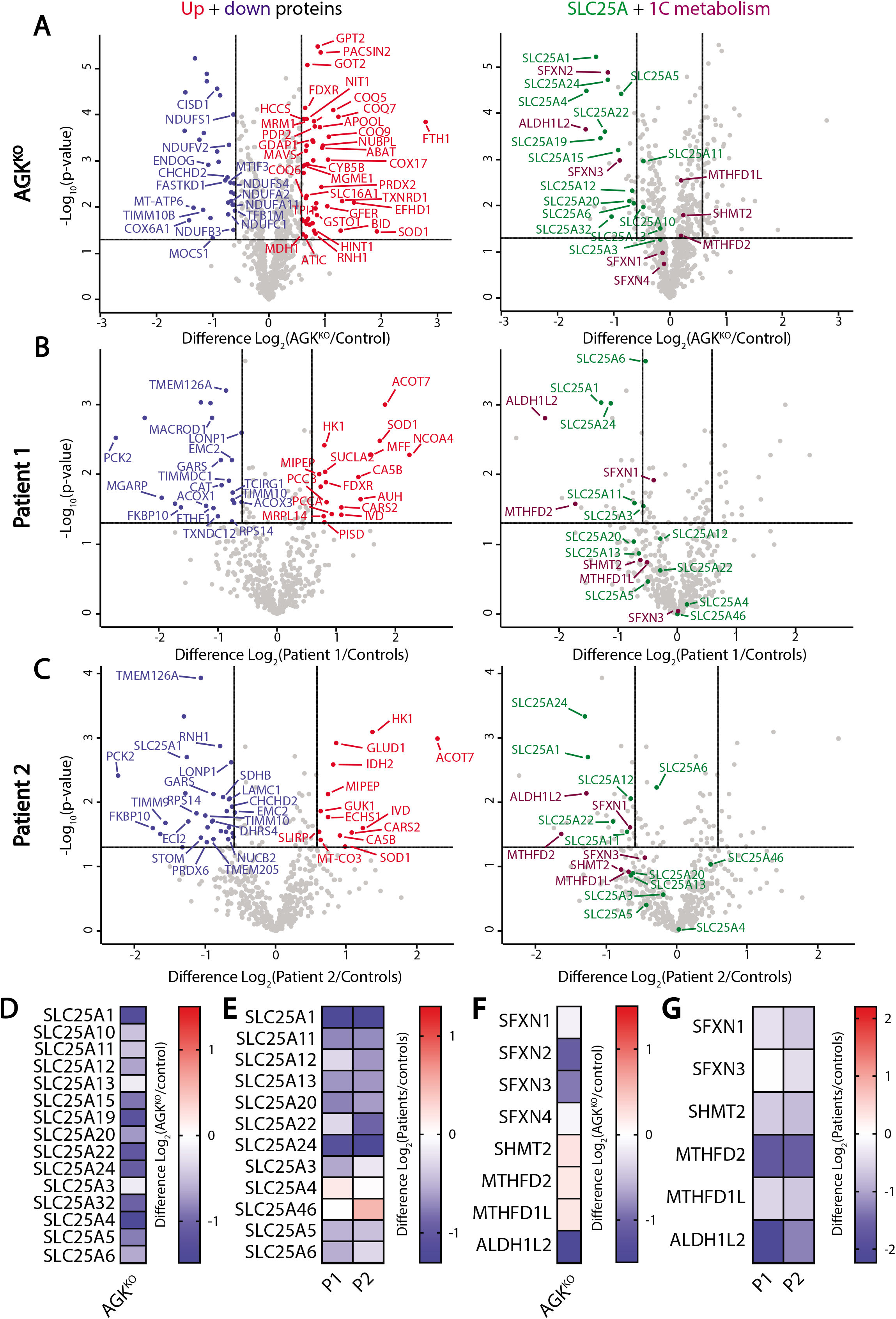
Mitochondrial 1C metabolism is remodelled in Sengers syndrome. **(A)** Mitochondria were isolated from AGK_KO_ and AGK_KO_ cells re-expressing AGK_WT 3×FLAG_ and subjected to label-free quantitative mass spectrometric analysis. Volcano plots depict the relative levels of proteins in AGK_KO_ mitochondria compared to control HEK293. n=3 biological replicates. Significantly altered proteins are located outside the lines (p-value: <0.05). Up-regulated proteins (red), down-regulated (blue), SLC25A family members (green) and 1C enzymes (plum) are indicated. **(B and C)** Mitochondria isolated from three independent control fibroblast cell lines and two Sengers patient fibroblast cell lines (Patient 1 (P1) and Patient 2 (P2)) were subjected to label-free quantitative mass spectrometric analysis. Significantly altered proteins are located outside the lines (p-value: <0.05). Up-regulated proteins (red), down-regulated (blue), SLC25A family members (green) and 1C enzymes (purple) are indicated. **(D-G)** Log_2_ fold-change values (as compared to respective controls) were depicted for selected proteins in the fibroblasts and AGK_KO_ HEK cells.

### Sideroflexins are novel substrates of the TIM22 complex

To confirm that the reduction in SFXN levels in Sengers patient fibroblasts arise due TIM22 complex dysfunction, rather than loss of AGK lipid kinase activity, we analysed levels of SFXN proteins in other systems of TIM22 complex dysfunction. Firstly, we excluded changes in the abundance of SFXN proteins due to alterations in gene expression by measuring mRNA abundance for SFXN1, SFXN2, SFXN3 and MTHFD2. Expression of SFXN1, SFXN2 and SFXN3 was unchanged in AGK_KO_ HEK293 cells and Tim9_MUT_ HEK293 cells (Kang et al., 2019) (**Figure 2A**). Tim9 also functions in the TIM22 pathway (Kang et al., 2019), suggesting the observed changes to the SFXN proteins were indeed post-translational. Protein levels of SFXN1 and SFXN4 were reduced in AGK_KO_ HEK293 and Tim9_MUT_ HEK293 mitochondria, as confirmed by western blot (**Figure 2B**). On BN-PAGE, SFXN1 assembles into a complex migrating at ~132 kDa (**Figure 2C, lane 1**). The abundance of this complex was reduced in AGK_KO_ and Tim9_MUT_ mitochondria (**Figure 2C, lanes 2 & 3**), where the TIM22 complex is destabilised (**Figure 2C, lanes 6 & 7**); and the complex was absent in a *SFXN1* CRISPR/Cas9 genome-edited cell line generated in this study (**Figure 2C, lane 4; Supplemental Figure 1**).

**Figure 2.**
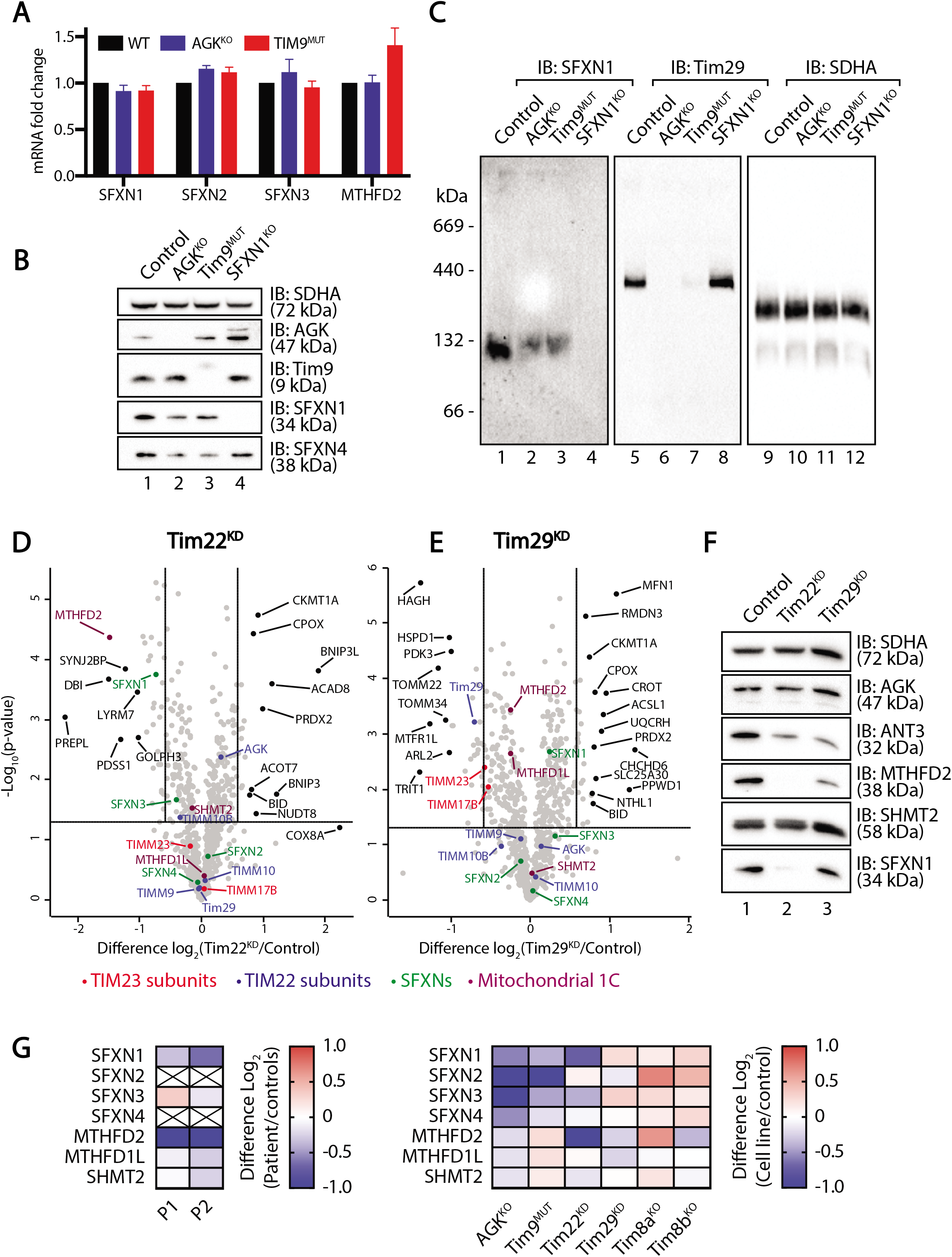
SFXN steady-state levels are reduced with TIM22 complex dysfunction. **(A)** Fold changes in mRNA expression for SFXN1, SFXN2, SFXN3 and MTHFD2 in AGK_KO_ and Tim9_MUT_ HEK293 compared to control HEK293 cells were quantified by RT-qPCR and are expressed as mean ± SD (n=3). **(B)** Mitochondrial lysates from control, AGK_KO_, Tim9_MUT_ and SFXN1_KO_ HEK293 cells were analysed by SDS-PAGE and western blotting with the indicated antibodies. **(C)** Mitochondria isolated from control, AGK_KO_, Tim9_MUT_ and SFXN1_KO_ HEK293 cells were solubilised in 1% digitonin containing buffer and analysed by BN-PAGE and immunoblotting with the indicated antibodies. **(D and E)** Mitochondria isolated from control HEK293 cells, **(D)** Tim22 KD, and (**E**) Tim29 knock-down (KD) cells were subjected to label-free quantitative mass spectrometric analysis. Volcano plots depict the relative levels of mitochondrial proteins in each sample compared to control. n=3 biological replicates. Horizontal cut-off represents p-value <0.05 and vertical cut-offs represent – and + 1.5x fold change. TIM23 complex subunits (red), TIM22 complex subunits (blue), SFXN proteins (green) and 1C metabolism enzymes (plum) are indicated. **(F)** Mitochondrial lysates from control, Tim22 KD and Tim29 KD HEK293 cells were analysed by SDS-PAGE and western blotting. **(G)** Log_2_ fold-change values (as compared to respective controls) were depicted for selected proteins in the indicated cell lines.

To further understand the requirement of the TIM22 complex for SFXN biogenesis, we performed label free quantitative mass spectrometry on mitochondria isolated from HEK293 cells depleted of either Tim22 or Tim29 via siRNA (**Figure 2D-E; Table 1**) and samples were retained for western blot analysis (**Figure 2F**). Tim22 is the central channel-forming unit of the TIM22 complex, while Tim29 maintains TIM22 complex integrity (Kang et al., 2016). In line with these different roles at the TIM22 complex, depletion of either Tim22 or Tim29 resulted in a distinct mitochondrial proteomic footprint. Tim22 knock-down substantially reduced the levels of SFXN1 and SFXN3 (**Figure 2D; Figure 2F-G**), but only had a minimal impact on the levels of SFXN2 and SFXN4 (**Figure 2D; Figure 2G**), perhaps due to the relatively short siRNA KD time course (72 hours). Interestingly, MTHFD2 was one of the most significantly downregulated proteins in the Tim22 KD (**Figure 2E-F, lane 2**), again suggesting that TIM22 complex dysfunction leads to remodelling of the mitochondrial 1C metabolism pathway. To the contrary, depletion of Tim29 led to reduced levels of Tim23 and Tim17b (**Figure 2E**), TIM22 substrates that are known to require Tim29 for efficient import (Callegari et al., 2016; Kang et al., 2016). Tim29 knock-down had no effect on the abundance of SFXN proteins (**Figure 2E-G**), as is the case for mitochondrial carrier proteins, which also use the TIM22 complex for import. In summary, the levels of SFXN proteins are reduced as a result of TIM22 complex dysfunction. This dependency is specific to the TIM22 complex rather than a non-specific response to stress, as Tim8a_KO_ and Tim8b_KO_ HEK293 mitochondria, which exhibit general mitochondrial dysfunction, but no TIM22 complex impairment (Kang et al., 2019), show no changes in abundance of SFXN proteins (**Figure 2G**).

As an additional biochemical measure to confirm that SFXNs are *bona fide* substrates of the TIM22 complex, we performed *in vitro* import and assembly assays. In vitro synthesised [_35_S]-SFXN1, [_35_S]-SFXN2, [_35_S]-SFXN3 and [_35_S]-Tim23 were incubated with mitochondria isolated from control and AGK_KO_ cells to allow for protein import and analysed by BN-PAGE to monitor protein assembly into complexes. At 60 minutes the assembly of SFXN2 and SFXN3 was significantly compromised in AGK_KO_ HEK293 mitochondria (**Figure 3A, quantified in 3C**), however no assembly defect was apparent for SFXN1 (**Figure 3A, lanes 1-6, quantified in 3C**). As expected, we observed no defects in the import of hTim23, which is known to require Tim29, but not AGK as an accessory receptor at TIM22 (Kang et al., 2016). The import defects closely mirrored the steady state levels of the SFXNs in AGK_KO_ mitochondria, where SFXN2 and SFXN3 were more substantially reduced than SFXN1 (**Figure 1A; Figure 1F**). The absence of a defect in SFXN1 import and assembly assessed through *in vitro* import could be explained by the fact that AGK is only a peripheral subunit of the TIM22 complex, and that the essential Tim22 pore is still present and functional in the absence of AGK. Indeed, more clear assembly defects were observed for SFXN1, SFXN2 and SFXN3 following import into Tim9_MUT_ cells (**Figure 3B-C**), which lack the essential intermembrane space chaperone Tim9. Together, these results suggest that SFXN1, SFXN2 and SFXN3 are novel substrates of the TIM22 complex and the import of SFXN proteins requires Tim9 and AGK, as well as Tim22 itself, while Tim29 appears to be dispensable.

**Figure 3.**
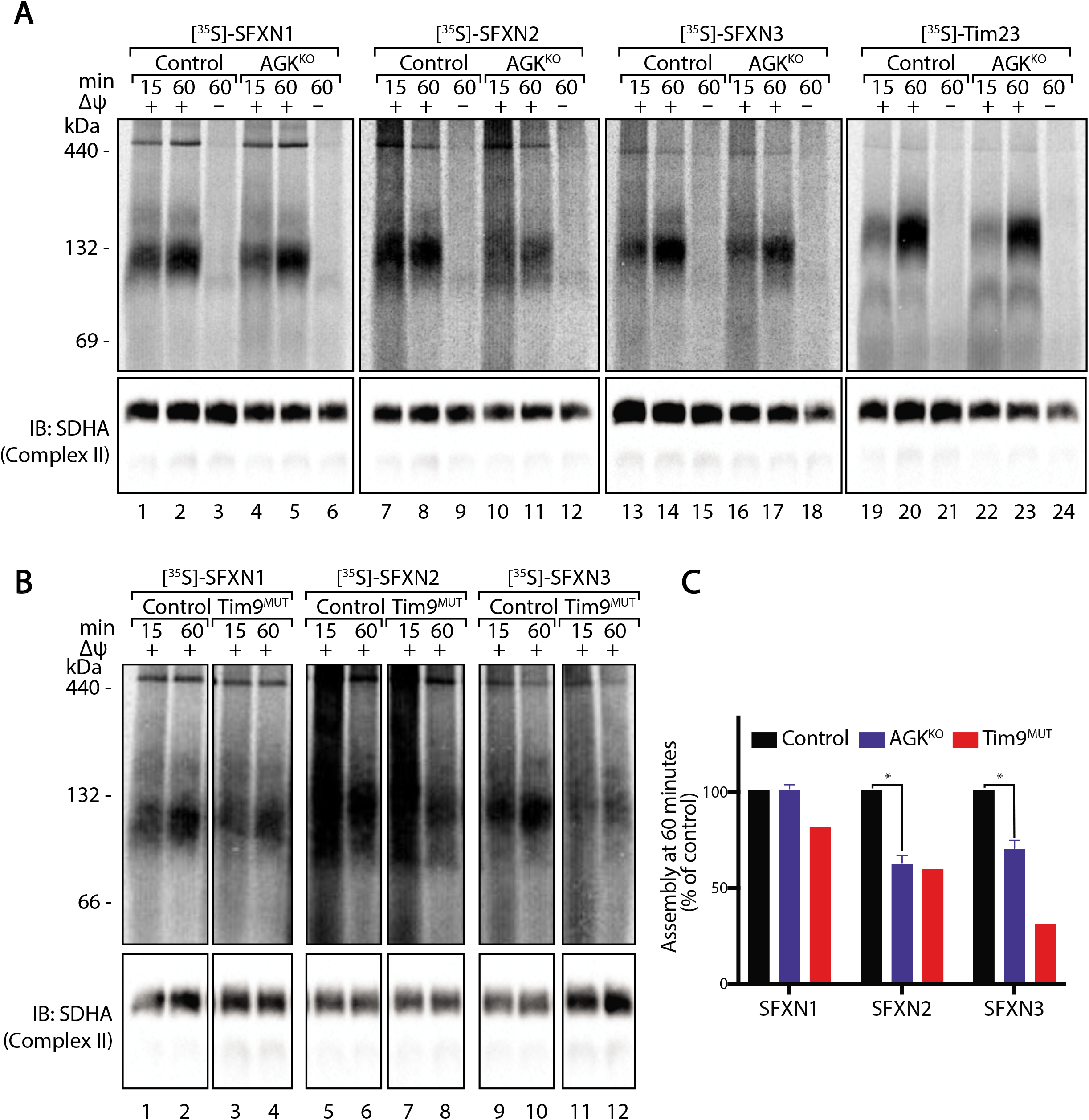
SFXNs are TIM22 complex substrates. **(A)** [_35_S]-SFXN1, [_35_S]-SFXN2, [_35_S]-SFXN3 or [_35_S]-Tim23 were incubated with mitochondria isolated from control HEK293 and AGK_KO_ cells for the indicated times in the presence or absence of a mitochondrial membrane potential (△ψ) prior to proteinase K (PK) treatment. Samples were solubilised in 1% digitonin containing buffer and analysed by BN-PAGE and autoradiography. Immunoblotting with SDHA was performed as a loading control. **(B)** [_35_S]-SFXN1, [_35_S]-SFXN2 or [_35_S]-SFXN3 were incubated with mitochondria isolated from control HEK293 and Tim9_MUT_ cells. Samples were solubilised in 1% digitonin containing buffer and analysed by BN-PAGE and autoradiography. Immunoblotting with SDHA was performed as a loading control. **(C)** Quantification of assembled protein at 60 minutes in control, AGK_KO_ and Tim9MU mitochondria. Graph depicts mean ± SD (n=3 for AGK_KO_, n=1 for Tim9_MUT_). Unpaired t-test, * p<0.05.

### Loss of AGK impairs growth in the absence of exogenous serine

We took note of the downward trend of mitochondrial 1C enzymes (MTHFD2 and SHMT2) in patient and model cell lines with TIM22 complex dysfunction (**Figure 1; Figure 2**). These enzymes are described as being localised to the mitochondrial matrix and are expected to utilise the TIM23 complex for import rather than the TIM22 complex. We therefore hypothesised that the reduction in the protein levels of MTHFD2 and SHMT2 was occurring due to a stress response induced downstream of reduced protein levels of SFXN proteins. To this end, we analysed our SFXN1_KO_ HEK293 cell line using label-free quantitative mitochondrial proteomics and confirmed that the absence of SFXN1 resulted in minor reductions in the levels of both MTHFD2 and SHMT2 (**Figure 4A-C**), suggesting that depletion of SFXN1 can partially underpin the remodelling of 1C metabolism observed following TIM22 complex dysfunction. Knock-out of SFXN1 also had no reciprocal effect on the abundance of TIM22 complex subunits (**Figure 4A, 4B & 4C**). Interestingly, depletion of SFXN1 also led to a reduction in the levels of SFXN2 and SFXN3 (**Figure 4A**). For SFXN2, but not SFXN3, this change was accompanied with a reduction in mRNA abundance (**Figure 4D**). The dependence of SFXN3 on SFXN1 for stability suggests that the two proteins may exist within a common complex. Together, this result suggests that depletion of SFXN levels due to TIM22 complex dysfunction may lead to further downstream remodelling of mitochondrial 1C metabolism. The mild effect observed in the SFXN1 KO may be explained by the presence of SFXN2 and SFXN3, which can perform redundant functions in serine transport (Kory et al., 2018).

**Figure 4.**
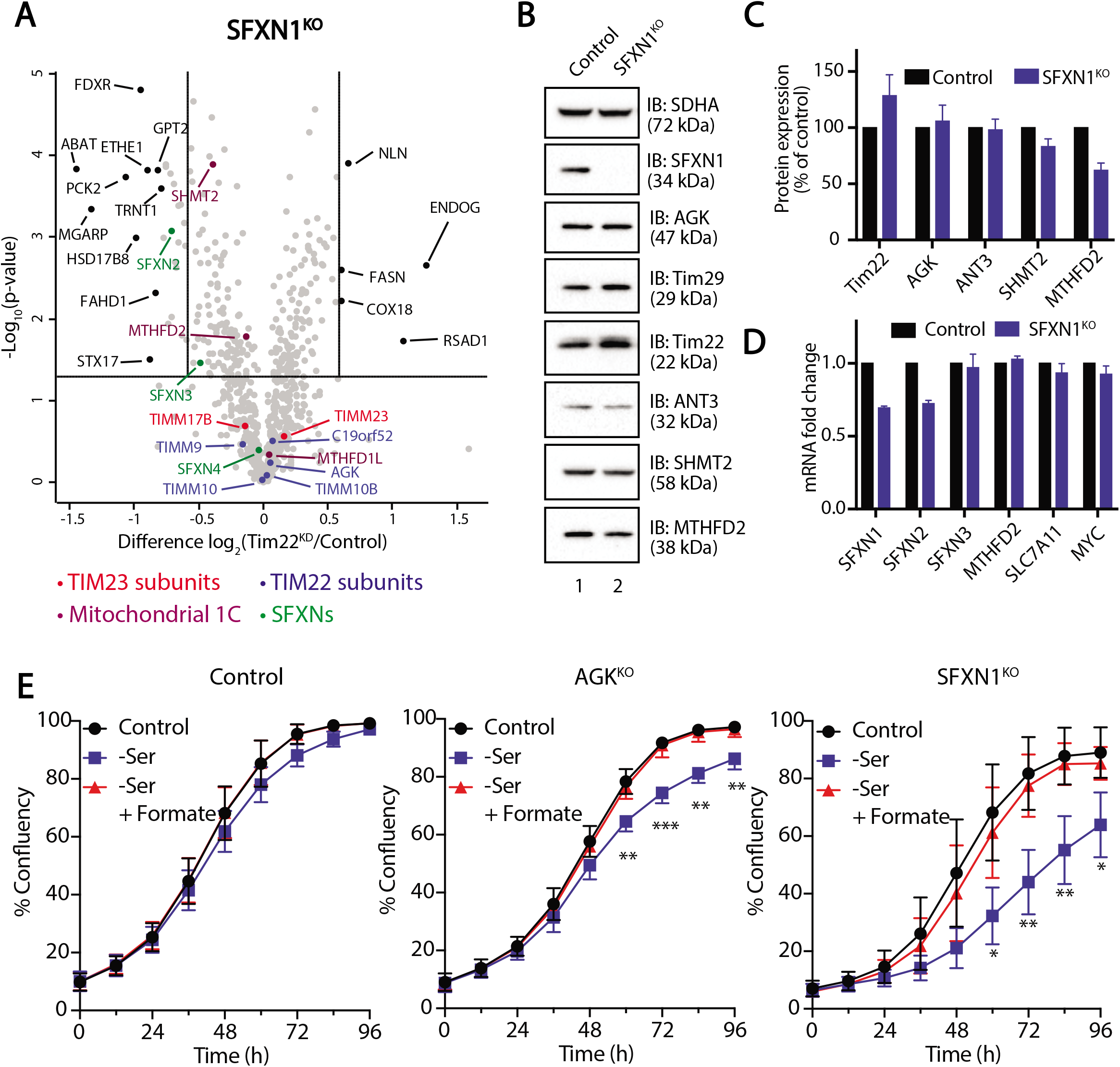
Loss of AGK limits cell proliferation in the absence of exogenous serine. **(A)** Mitochondria isolated from control and SFXN1_KO_ HEK293 cells were subjected to label-free quantitative mass spectrometric analysis. Volcano plots depict the relative levels of mitochondrial proteins in each sample compared to control. n= 3 biological replicates. Horizontal cut-off represents p=0.05 and vertical cut-offs represent – and + 1.5x fold change. TIM23 complex subunits (red), TIM22 complex subunits (blue), SFXN proteins (green) and 1C metabolism enzymes (plum) are indicated. **(B)** Mitochondrial lysates from control and SFXN1_KO_ HEK293 cells were analysed by SDS-PAGE and western blotting. (**C**) Relative protein levels of selected proteins were quantified and are represented as the mean ± SD (n=3). **(D)** Relative fold changes of mRNA expression for SFXN1, SFXN2, SFXN3, MTHFD2, SLC7A11, and MYC in control, and SFXN1_KO_ HEK293 cells were quantified and are represented as the mean +/− SD (n=3). **(E)** Proliferation of control, AGK_KO_ and SFXN1 KO HEK293 cells was monitored in complete media, serine-free media, and serine-free media supplemented with 1 mM formate. Confluency was measured at 12-hour intervals and was depicted as mean ± SD, n=4. Unpaired t-test, * p<0.05, ** p<0.01, *** p<0.001.

To confirm that reduced SFXN levels downstream of AGK/TIM22 complex dysfunction result in a functional defect in serine utilisation at the mitochondrion, we monitored cell proliferation of control and AGK_KO_ cells in serine-free media (**Figure 4E**). Under these conditions, intracellular serine levels are reduced and can only be supplied in limited quantities through the *de novo* serine biosynthesis pathway. Normal proliferation under these conditions requires efficient mitochondrial 1C metabolism and any defects in this pathway, including reduced import of serine into the mitochondria, are likely to limit nucleotide synthesis and manifest as a growth defect. Indeed, while AGK_KO_ HEK293 cells have no proliferation defect under standard culturing conditions (Kang et al., 2017) growth in serine-free media resulted in a mild but significant proliferation defect (**Figure 4E, middle panel**). This defect could be rescued through supplementation with 1 mM formate, a key product of the mitochondrial 1C cycle. SFXN1_KO_ cells were also rescued by supplementation with formate (**Figure 4E, right panel**). Together, these results suggest that AGK/TIM22 complex dysfunction reduces the efficiency of metabolic flux through the mitochondrial arm of the 1C metabolism pathway, most likely by reducing the efficiency of SFXN import and assembly into the inner membrane.

## Discussion

In this study, we set out to characterise the mitochondrial proteome of Sengers syndrome patient fibroblasts with a view to identifying altered pathways following loss of AGK, a subunit of the metazoan TIM22 complex. As expected, our analysis demonstrated reduced abundance of SLC25A proteins, the prototypical substrates of the TIM22 complex. The biogenesis of SLC25A members has not been studied systematically, and our observation that steady state abundance of all detected SLC25A members was reduced in AGK_KO_ HEK293 mitochondria suggests that most members of this protein family relies on AGK and the TIM22 complex for efficient biogenesis.

The proteomic approach employed in this study also uncovered extensive remodelling of key players involved in mitochondrial 1C metabolism in fibroblasts from patients with Sengers syndrome, including reduced levels of MTHFD2, MTHFD1L, SHMT2, ALDH1L2 and the SFXN proteins. SFXN proteins are inner mitochondrial membrane serine transporters that are required for mitochondrial 1C metabolism (Kory et al., 2018). Based on our analysis showing reduced import and assembly of SFXN proteins in mitochondria lacking AGK and Tim9, we suggest that that this family of proteins represent novel substrates of the TIM22 complex. Indeed, independent mass spectrometric analysis of AGK_KO_ HEK293 cells shows a striking downregulation of SFXN proteins and identified SFXN2 as a high confidence client protein of AGK (Vukotic et al., 2017), while SFXN1 has also been enriched following Tim22 immunoprecipitation (Callegari et al., 2016). The identification of SFXNs as TIM22 complex substrates prompts a re-think of the mechanisms of TIM22 complex membrane insertion. The two previously known substrate classes possess 4 or 6 transmembrane domains and were thought to be inserted as a series of hairpin loops (Rehling et al., 2003). SFXN proteins possess 5 transmembrane domains and are not compatible with this model of membrane insertion. Further biochemical analysis will be required to dissect the steps involved in their biogenesis.

Unlike the SFXN proteins, ALDH1L2, SHMT2 and MTHFD2 are thought to be soluble matrix proteins and are presumably not substrates of the TIM22 complex. Consistent with this is the downregulation or turnover of SHMT2 and MTHFD2 in SFXN1_KO_ cells (**Figure 4A**). These enzymes have been shown to be upregulated by ATF4 in models of mitochondrial disease associated with mtDNA mutation (Bao et al., 2016; Khan et al., 2017; Nikkanen et al., 2016), although we did not explore the possibility of an ATF4 response in Sengers patient cells. Mitochondrial 1C metabolism plays a crucial role in redox balance through generation of glycine for glutathione synthesis (Ducker and Rabinowitz, 2017) and NADPH for glutathione recycling (Fan et al., 2014). It is therefore possible that this response limits that ability of the cell to combat oxidative stress that occurs during mitochondrial dysfunction. It is also important to consider the contribution of serine catabolism to mitochondrial translation, as serine contributes to the formylation of Met-tRNAfMet and perturbations in this pathway result in OXPHOS defects (Minton et al., 2018). This is apparent in our AGK_KO_ proteomics data (**Table 1**) where there is a striking defect in the levels of numerous mtDNA encoded subunits, including MT-ATP6, MT-ND1, and MT-ND4. This research naturally leads to the idea that supplementation with formate might benefit patients with Sengers syndrome, or other mitochondrial diseases with perturbations to 1C metabolism. While folate deficiency has been discussed in the context of mitochondrial disease (Garcia-Cazorla et al., 2008; Ormazabal et al., 2015), the underlying mechanisms are poorly understood. One-carbon metabolism is known to play a crucial role in development (Momb et al., 2013), but its function in adult tissues is not clear (Ducker and Rabinowitz, 2017). It is possible that under conditions of mitochondrial dysfunction and oxidative stress, one-carbon metabolism plays an important anti-oxidant role.

The identification of SFXNs as substrates of the TIM22 complex positions the TIM22 complex as an indirect regulator of 1C metabolism, as SFXN proteins determine the amount of mitochondrial serine that is available to SHMT2. It is tempting to speculate that the levels of the TIM22 complex could be modulated to regulate the level of SFXNs in the inner membrane and thus determine flux through the mitochondrial 1C pathway. Interestingly, components of the mitochondrial 1C pathway are overexpressed in cancers (Nilsson et al., 2014; Rosenzweig et al., 2018). AGK is also overexpressed in several cancer types (Bektas et al., 2005; Chen et al., 2013; Wang et al., 2014), although its contribution to tumorigenesis has been thought to relate to its lipid kinase activity. These results raise the hypothesis wherein overexpression of components of the TIM22 complex in cancer might serve a need for increased biogenesis of SFXN proteins.

Overall, the results presented here identify SFXNs as novel substrates of the TIM22 complex and mitochondrial 1C metabolism as a pathway that is dysregulated in Sengers syndrome. The loss of SFXNs in Sengers syndrome might lead to downstream remodelling of 1C metabolism that is potentially maladaptive. Supplementation with formate could represent a novel therapeutic strategy, although the requirements for 1C metabolism in non-proliferative tissues are not clear. The TIM22 complex is positioned as a potential regulator of mitochondrial 1C metabolism in both health and pathologies such as cancer and mitochondrial disease.

## Acknowledgements

T.J. and K.M.F. are supported by Australian Government Research Training Program (RTP) Scholarships. D.H.H is supported by a Melbourne International Research Scholarship. T.J and D.H.H are supported by Mito Foundation PhD Top-Up scholarships. D.S. is supported by the Research Fellowship from the Mito Foundation. We acknowledge funding from the Australian Research Council (DP170101249 to D.S) and National Health and Medical Research Council) and National Health and Medical Research Council (NHMRC Project Grant 1140906 to D.A.S; NHMRC Fellowship 1140851 to D.A.S; NHMRC Fellowship 1155244 to D.R.T). N.J.C. is supported by a Fellowship (MCRF16002) from the Victorian Government Department of Health and Human Services acting through the Victorian Cancer Agency.

## Author contributions

Conceptualization, T.J. and D.S.; Methodology, T.J., D.H.H., C.P., Y.K., D.A.S., D.S.; Formal Analysis, T.J., D.H.H, K.M.F., D.A.S; Investigation, T.J., D.H., K.M.F., Y.K., C.P.; Resources, N.J.C., D.T., D.A.S., D.S.; Writing – Original Draft, T.J and D.S.; Writing – Review & Editing, all authors; Visualisation, T.J., D.H., D.S., Supervision, N.J.C., D.T., D.A.S., D.S., Project Administration, D.S., Funding Acquisition, D.S.

## Methods

### Cell lines, cell culture and siRNA transfection

Flp-In_TM_ T-Rex_TM_ 293 (Thermo Fisher Scientific) and primary patient fibroblasts (Calvo et al., 2012) were cultured in Dulbecco’s modified Eagle’s medium (DMEM; Thermo Fisher Scientific), containing 1% [v/v] penicillin-streptomycin (Thermo Fisher Scientific) and supplemented with 5% [v/v] foetal bovine serum (Sigma). siRNA transfection was performed in cells plated overnight using scrambled siRNA (Sigma) or siRNA targeting Tim22 (5’ CCAUUGUGGGAGCCAUGUU 3’) (Sigma) or Tim29 (5’ GGCUCUUCGAUGAGAAGUA 3’) (Sigma). Briefly, siRNA was transfected at 10 nM using DharmaFECT (Dharmacon) according to the manufacturer’s instructions. Cells were transfected a second time 48 hours after the first transfection and harvested 72 hours after the first transfection.

### Gene editing and screening

Editing of the *SFXN1* gene was carried out using the pSpCas9(BB)-2A-GFP CRISPR-Cas9 construct (a gift from F. Zhang; Addgene) (Ran et al., 2013). Guide RNAs targeting exon 7 (coding exon 6) of the SFXN1 gene were designed using CHOPCHOP. An oligonucleotide duplex formed from (5’ CACCGCGTTCGCCGACTCCCCCAAG 3’ and 5’ AAACCTTGGGGGAGTCGGCGAACGC 3) was ligated into pSpCas9(BB)-2A-GFP and transfected into Flp-In_TM_ T-Rex_TM_ 293 cells, and single cells were obtained via FACS based on GFP fluorescence. Sorted cells were allowed to expand prior to screening. Screening was performed through western blotting with a SFXN1 antibody and clones were genetically verified using genomic sequencing (Supplemental Figure 1). Mass spectrometry also validated the absence of SFXN1 protein in the knockout cell line.

### Quantitative RT-PCR

Following RNA extraction using a NucleoSpin RNA kit (Macherey-Negal), cDNA was synthesised with the Transcriptor First Strand cDNA Synthesis kit (Roche). Gene expression was determined using SYBR-green qPCR on the Lightcycler 480 (Roche). Gene expression was normalised to GAPDH and ACTB and determined using the △△Ct method. Primer sequences used in this study were:

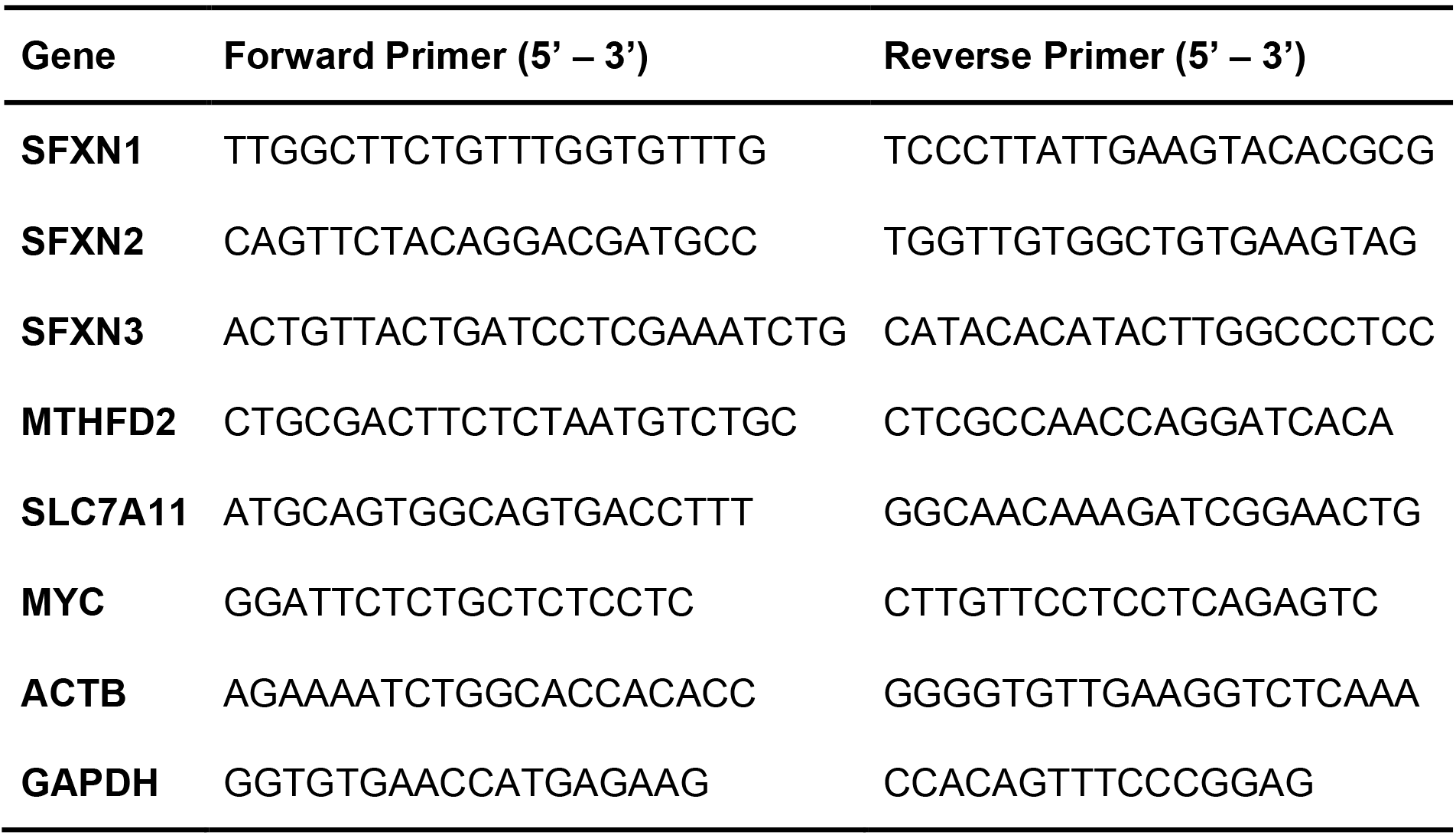

### Mitochondrial isolation, gel electrophoresis and immunoblot analysis

Mitochondria were isolated from cultured mammalian cells through differential centrifugation as described previously (Kang et al., 2017). Briefly, cultured cells were harvested in Phosphate Buffered Saline (PBS) and isolated by centrifugation at 500 *g*. Cells were homogenised in isolation buffer (20 mM HEPES-KOH (pH 7.6), 220 mM mannitol, 70 mM sucrose, 1 mM EDTA, 0.5 mM PMSF and 2 mg/mL BSA) and the lysate was centrifuged at 800 *g* to remove nuclear debris and intact cells. The supernatant containing mitochondria was centrifuged at 12,000 *g* to obtain a crude mitochondrial pellet. Protein concentration in the mitochondrial pellet was determined using the Pierce BCA protein assay kit (Thermo Fisher Scientific).

Tris-Tricine SDS-PAGE was performed as described previously (Kang et al., 2017). Solutions containing 10 or 16% [v/v] acrylamide solution (49.5% acrylamide, 1.5% bis-acyrlamide) were made up in tricine gel buffer (1M Tris-Cl, 0.1% [w/v] SDS, pH 8.45; 13% [v/v] glycerol included in the 16% mix). These solutions were used to pour 10-16% gels using a gradient mixer. Following polymerisation, a stacking gel (4% [v/v] acrylamide solution in tricine gel buffer) was overlaid onto the gradient gel. Polymerisation of both the gradient and stacking gels was achieved through sequential addition of TEMED (Sigma) and 10% APS (Sigma). Electrophoresis was performed using Tris-Tricine SDS-PAGE anode buffer (50 mM Bis-Tris, pH 7.0) and cathode buffer (0.1 M Tris, 0.1 M Tricine, 0.1% [w/v] SDS, pH 8.45). Pelleted mitochondria to be analysed were resuspended in SDS loading dye (200 mM Tris-Cl (pH 6.8), 0.4 M DTT, 8% [w/v] SDS, 40 % [v/v] glycerol, 0.2 % bromophenol blue) and boiled.

Blue-Native (BN) PAGE was performed as described previously (Kang et al., 2017). Solutions containing 4 or 16% [v/v] acrylamide solution in BN gel buffer (66 mM ε-amino n-caproic acid, 50 mM Bis-Tris, pH 7.0) were used to pour 4-16% gradient gels. Following polymerisation, a stacking gel (4% [v/v] acrylamide solution in BN gel buffer) was overlaid onto the gradient gel. Polymerisation was initiated through sequential addition of TEMED and 10% APS. Electrophoresis was carried out overnight at 4 °C using BN anode (50 mM Bis-Tris, pH 7.0) and cathode buffer (50 mM Tricine, 15 mM Bis-Tris, 0.02% [w/v] Coomassie blue G250). Pelleted mitochondria to be analysed were solubilised in digitonin solubilisation buffer (20 mM Bis-Tris, 50 mM NaCl, 10% [v/v] glycerol, pH 7.4, 1% [w/v] digitonin). BN loading dye (0.5% [w/v] Coomassie blue G250, 50 mM ε-amino n-caproic acid, 10 mM Bis-Tris, pH 7.0) was added to the clarified supernatant prior to loading.

Following electrophoresis, gels were transferred onto PVDF membranes (0.45 μM Immobilon-P, Merck) using the Owl HEP-1 Semidry Electroblotting system (Thermo Fisher Scientific). Following incubation with primary antibodies, horseradish peroxidase coupled secondary antibodies (Sigma) and Clarity Western ECL Substrate (BioRad) were used for detection. Images were obtained using the ChemiDoc XRS+ imaging machine (BioRad).

### *In vitro* protein import and autoradiography

Open reading frames encoding SFXN1, SFXN2, SFXN3 or Tim23 were cloned into pGEM4z and used for transcription with the mMESSAGE mMACHINE SP6 kit (Thermo Fisher Scientific) according to the manufacturer’s instructions. Radiolabelled protein was translated from mRNA using the rabbit reticulocyte lysate system (Promega) and _35_S-labelled methionine, according to the manufacturer’s instructions. Isolated mitochondria were resuspended in mitochondrial import buffer (250 mM sucrose, 5 mM magnesium acetate, 80 mM potassium acetate, 10 mM sodium succinate, 1 mM DTT, 5 mM ATP, 20 mM HEPES-KOH, pH 7.4), and imports were performed through incubation with radiolabelled proteins for the desired incubation time at 37 °C, and in the presence or absence of 10 μM FCCP (to dissipate membrane potential). Following import, each reaction was treated consecutively with Proteinase K (50 μg/mL for 10 minutes on ice) and PMSF (1 mM for 5 minutes on ice) prior to re-isolation for SDS-PAGE and BN-PAGE analysis. Radioactive signals were detected using a Typhoon phosphorimager (GE healthcare). Analysis of autoradiography was performed using ImageJ software to calculate the intensity of each band. Background intensities were calculated by averaging the intensity of multiple areas of the gel away from the bands.

### Quantitative mass spectrometry and data analysis

A total of 50 μg of protein from whole-cell or isolated mitochondria were estimated using a Pierce BCA Assay Kit (Thermo Fisher Scientific). Pellets were solubilised in solubilisation buffer (1 % [w/v] SDC, 100 mM Tris pH 8.1, 40 mM chloroacetamide (Sigma) and 10 mM tris(2-carboxyethylphosphine hydrochloride (TCEP; BondBreaker, Thermo Fisher Scientific) for 5 minutes at 99 °C with 1500 rpm shaking followed by 15 minutes sonication in a water bath sonicator. Proteins were digested with trypsin (Thermo Fisher Scientific) at a 1:50 trypsin:protein ratio at 37 °C overnight. The supernatant was transferred to stagetips containing 3×14G plugs of 3_MT_MEmpore_TM_ SDB-RPS substrate (Sigma) as described previously (Kulak et al., 2014; Stroud et al., 2016). Ethyl acetate or isopropanol 99% [v/v] and 1% TFA [v/v] was added to the tip before centrifugation at 3000 *g* at room temperature. Stagetips were washed first with ethyl acetate or isopropanol (99% [v/v]) and TFA (1% [v/v]) solution and then subjected to a second wash containing 0.2 % [v/v] TFA. Peptides were eluted in 80% [v/v] acetonitrile and 1% [w/v] NH_4_OH and acidified to a final concentration of 1% [v/v] TFA prior to during in a CentriVap Benchtop Vacuum Concentrator (Labconco). Peptides were reconstituted in 0.1% TFA and 2% CAN for analysis by liquid chromatography (LC) – MS/MS.

LC MS/MS was carried out on a QExactive plus Orbitrap mass spectrometer (Thermo Fisher Scientific) with a nanoESI interface in conjunction with an Ultimate 3000 RSLC nanoHPLC (Dionex Ultimate 3000. The LC system was equipped with an Acclaim Prepmap nano-trap column (Dionex C18, 100 Å, 75 μM x 50 cm). The tryptic peptides were injected to the enrichment column at an isocratic flow of 5 μL/min of 2% [v/v] CH_3_CN containing 0.1% [v/v] formic acid for 5 min applied before the enrichment column was switched in-line with the analytical column. The eluents were 5% DMSO in 0.1% [v/v] formic acid (solvent A) and 5% DMSO in 100% [v/v] CH_3_CN and 0.1% [v/v] formic acid (solvent B). The flow gradient was (i) 0-6 min at 3% B, (ii) 6-95 min at 3-22% B, (iii) 95-105 min at 22-40% B, (iv) 105-110 min at 40-80% B, (v) 110-115 min at 80% B, (vi) 115-117 min at 80-3% B. Equilibration was performed with 3% B for 10 minutes before the next sample injection. The QExactive plus mass spectrometer was operated in the data-dependent mode. Full MS1 spectra were acquired in positive mode, 70000 resolution, AGC target of 3e_6_ and maximum IT time of 50 ms. A loop count of 15 on the most intense targeted peptide were isolated for MS/MS. The isolation window was set to 1.2 m/z and precursors fragmented using stepped normalised collision energy of 28, 30 and 32. MS2 resolution was at 17500, AGC target at 2e6 and maximum IT time of 50 ms. Dynamic exclusion was set to be 30 s.

Raw files were processed using the MaxQuant platform (version 1.6.5.0) (Cox and Mann, 2008) and searched against the UniProt human database (June 2019) using default settings for an LFQ experiment with match between runs enabled. The proteinGroups.txt output from the search was processed in Perseus (version 1.6.2.2) (Tyanova et al., 2016). Briefly, entries “Only identified by site”, “Reverse” and “Potential contaminant” were removed from the data-sets. Log_2_ transformed LFQ intensities were grouped (control, knockout, patient) according to each experiment and filtered to have 2 out of 3 valid values in each group. Isolated mitochondria experiments were annotated for proteins present in the Mitocarta2.0 (Calvo et al., 2016) through matching by gene name. Mitocarta2.0 positive rows were filtered to include only mitochondrial entries and normalised using the “Subtract row cluster” function with “Known mitochondrial” entries from the IMPI (2017) (Smith and Robinson, 2016) database as reference. Two samples t-tests were performed between groups using p-value for truncation (threshold p-value = 0.05). Volcano plots were generated via scatter plots by selecting “Student’s T-test difference” and “-Log Student’s T-test p-value”.

### Cell proliferation measurement

Confluency of cells in 96-well plates was tracked over 96 hours using IncuCyte FLR (Essen BioSciences) following manufacturer’s guidelines. 5000 cells were plated in indicated media and allowed to adhere 2 hours prior to the first reading.

---

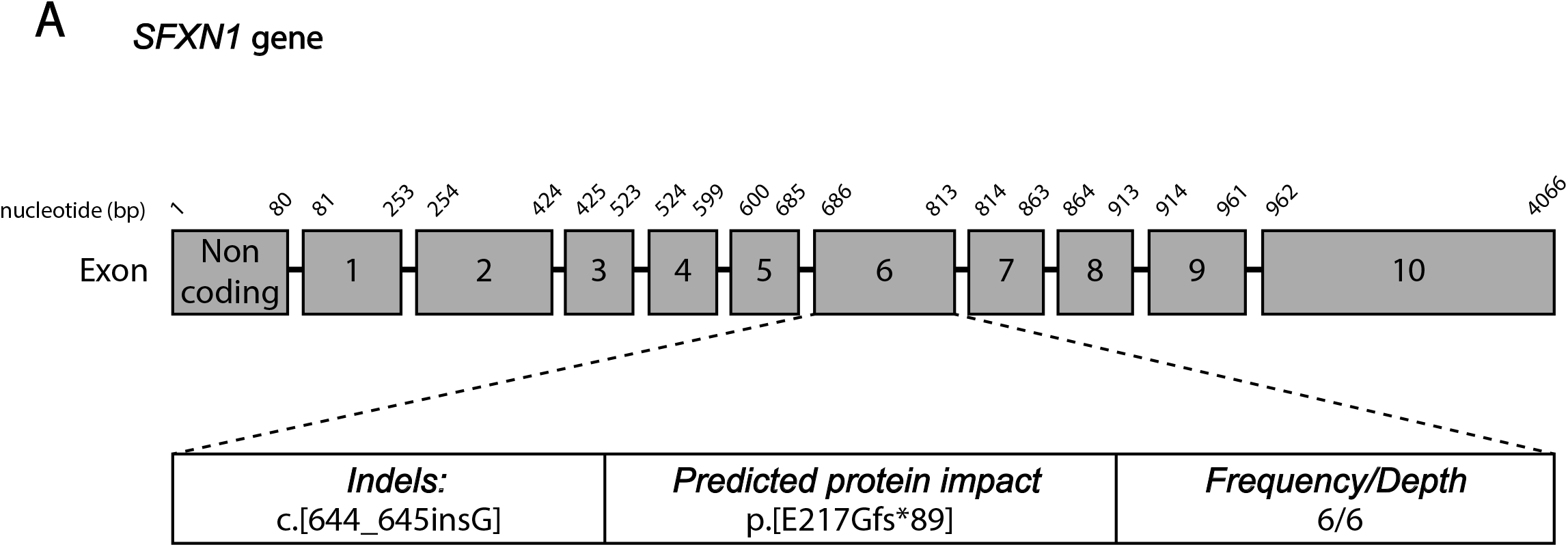

